# Single cell transcriptomic analyses of human heart failure with preserved ejection fraction

**DOI:** 10.1101/2025.03.06.641949

**Authors:** Virginia S. Hahn, Mark Chaffin, Bridget Simonson, Sydney C. Jenkin, Abigail S. Mulligan, Kenneth C. Bedi, Kenneth B. Margulies, Carla A Klattenhoff, Kavita Sharma, David A. Kass, Patrick T. Ellinor

**Author notes:** **Address for Correspondence:** Patrick T. Ellinor, MD, PhD, Heart and Vascular Institute, Mass General Brigham, 55 Fruit Street, GRB810, Boston, MA 02214. These authors contributed equally (Virginia S. Hahn, Mark Chaffin).

## Abstract

**Background:** Heart failure with preserved ejection fraction (HFpEF) is a poorly understood, multi-system disease with high morbidity and mortality. To improve our understanding of its underlying biology, we used single-nucleus RNA sequencing (snRNA-seq) to characterize cell-specific gene expression patterns in human HFpEF myocardium.

**Methods:** Septal myocardial biopsies (2-3 mg) from 30 HFpEF patients and 29 non-failing donor controls were analyzed using the 10X Genomics platform, with nuclei isolated from combined samples (6 patients/pool). Genotype-based demultiplexing was performed with *souporcell*, and gene expression quantified with *CellRanger* and *CellBender*. After quality control, nuclei were clustered and annotated by cell types based on specific marker genes. Differential expression (DE) by cell-type in HFpEF vs controls was performed using *limma-voom* and functional analysis performed using Gene Set Enrichment Analysis. Data were compared to dilated cardiomyopathy (DCM) using prior snRNA-seq in DCM vs respective controls.

**Results:** We successfully demultiplexed pooled myocardial biopsies, assigning >75% of droplets to individual patients. From eight pooled samples (19 HFpEF, 24 controls), we recovered 48,886 nuclei and identified 14 cell types. Cardiomyocytes (5159 differentially expressed [DE] genes, 36%) and fibroblasts (5905 DE genes, 49%) showed the most DE genes, while endothelial cells (2143), pericytes (1812), and macrophages (1405) had fewer. Enriched pathways common to multiple cell types included transcription/translation, immune activation, metabolism, and protein quality control. Of 7848 DE genes identified via pseudo-bulk snRNA-seq, 51% were DE in fibroblasts and 47% in cardiomyocytes, compared to <20% in other cell types. Unlike dilated cardiomyopathy (DCM), sub-clustering fibroblasts did not reveal an activated fibroblast population in HFpEF. Comparative analysis between HFpEF and DCM identified transcriptional differences primarily in cardiomyocytes.

**Conclusions:** This study demonstrates the power of genotype-based demultiplexing for single-cell transcriptomic analyses of small endomyocardial biopsies and identifies cardiomyocytes as the principal cell type with distinct transcriptional changes in HFpEF versus DCM. These findings, coupled with differential gene expression and functional pathway analyses, illuminate HFpEF pathways and may nominate compelling targets for future mechanistic studies and therapeutic efforts for HFpEF.

**Clinical Perspective:** **What is new?**

We successfully used genotype-based demultiplexing to perform single nucleus RNA-seq from myocardial biopsies.

The snRNA-seq analysis revealed distinct enrichment of pathways related to transcription/translation, immune activation, metabolism, and protein quality control across multiple cell types in HFpEF.

In contrast to DCM, HFpEF is distinguished by the absence of an activated fibroblast population and a predominance of transcriptional differences within cardiomyocytes, highlighting a distinct disease mechanism.

**What are the clinical implications?**

The cell-selective HFpEF myocardial transcriptional landscape highlights altered metabolism, protein translation and quality control, immune activation and growth/matrix organization.

These pathways provide compelling new targets for developing effective treatments.

## INTRODUCTION

Heart failure affects nearly 7 million adults in the United States alone^1^, imposing over $30 billion in health-care costs per year.^2^ Worldwide, the incidence approaches 65 million.^3^ Approximately half have heart failure with preserved ejection fraction (HFpEF), a syndrome where resting systolic function appears normal, yet patients exhibit similar symptoms and outcomes as those with reduced systolic function. ^4^ While long viewed as a disorder of diastolic function, HFpEF has become dominated by an obesity cardiometabolic syndrome phenotype, impacting multiple organs and cell types including cardiomyocytes^5^, vascular^6^, immune^7^, and interstitial cells^8^. Despite a normal resting EF, cardiac insufficiency particularly under conditions of exertional demand is observed. Current HFpEF animal models often incorporate both metabolic and hemodynamic stressors (e.g., two-hit models), but still fail to fully recapitulate the complex, multi-system pathophysiology observed in human HFpEF. Critically, limited molecular and cellular data from human myocardium has hindered validation of such preclinical models and identification of high-value therapeutic targets for HFpEF. This underscores the vital importance of human tissue analysis in elucidating the myocardial mechanisms driving this disease.

Our laboratory has previously reported on human HFpEF myocardial transcriptomics using bulk RNA-seq^9^, myocardial metabolomics^10^, myocardial proteomics ^11^, integrated analysis of glycolysis^12^, ultrastructural analysis^13^, and myocyte sarcomere function^5^. Collectively, these studies revealed a potent impact of marked obesity often present in HFpEF patients, with signatures of depressed fat and glucose metabolism, altered protein translation and quality control, lipid accumulation, and immune activation. A major unresolved question is how these abnormalities segregate among different myocardial cell types, analysis recently applied to a variety of other cardiac disease syndromes^14–18^. This is invaluable for identifying key signatures pertinent to cardiomyocytes, and equally to cell-types of lower abundance and/or transcriptional activity, such as immune, matrix, and vascular cells. Systemic inflammation is found in serum proteomics of human HFpEF, ^20–22^ but its relation to immune cell changes in the myocardium remains unknown. In this regard, single-nucleus RNA sequencing (snRNA-seq) studies of human myocardium from patients with dilated cardiomyopathy (DCM) and hypertrophic cardiomyopathy (HCM) have identified a distinct activated fibroblast population^14^, and altered proportions of immune cell subclusters with greater diversification in different disease groups.^14,15^ Whether or how this pertains to human HFpEF is also unknown.

To define cell-specific signatures in human HFpEF myocardium, we employed snRNA-seq. This was used as opposed to single-cell RNA sequencing given the large size and rod-like shape of cardiomyocytes that limits their passage through the microfluidic devices used to perform such analysis. Endomyocardial biopsies from a well-phenotyped HFpEF cohort were compared to ventricular myocardium obtained from non-failing (NF) donor controls using a combination of differential gene expression with functional annotation and sub-cluster analysis. Comparisons were also made to a prior snRNA-seq analysis of human DCM and respective controls.^14^

## METHODS

### Data Availability

De-identified raw sequence data are available in dbGaP with controlled-access. Summary level data are available in the Single Cell Portal at the Broad Institute.

### Study Population and Study Design

**Figure 1** provides a schematic overview of the study. This was a cross-sectional study using myocardial tissue samples from two biobanks. Patients presenting to the JHU HFpEF clinic were referred for clinically indicated right heart catheterization with research endomyocardial biopsy. HFpEF diagnosis was based on established consensus criteria^4^, including right heart catheterization demonstrated elevation of pulmonary capillary wedge pressure at rest or with supine exercise as described previously^9^. Detailed inclusion and exclusion criteria are in the Supplement. Ventricular septal endomyocardial biopsies and location-matched non-failing (NF) control tissue from organ donor subjects were obtained as described^19^, frozen in liquid nitrogen, and stored in a -160° freezer. The study was approved by the Johns Hopkins University Institutional Review Board (HFpEF) and the Perelman School of Medicine Institutional Review Board (controls). Demographics and clinical characteristics between HFpEF and controls were compared by Wilcoxon tests (continuous) or Fisher’s exact test (categorical).

**Figure 1.**
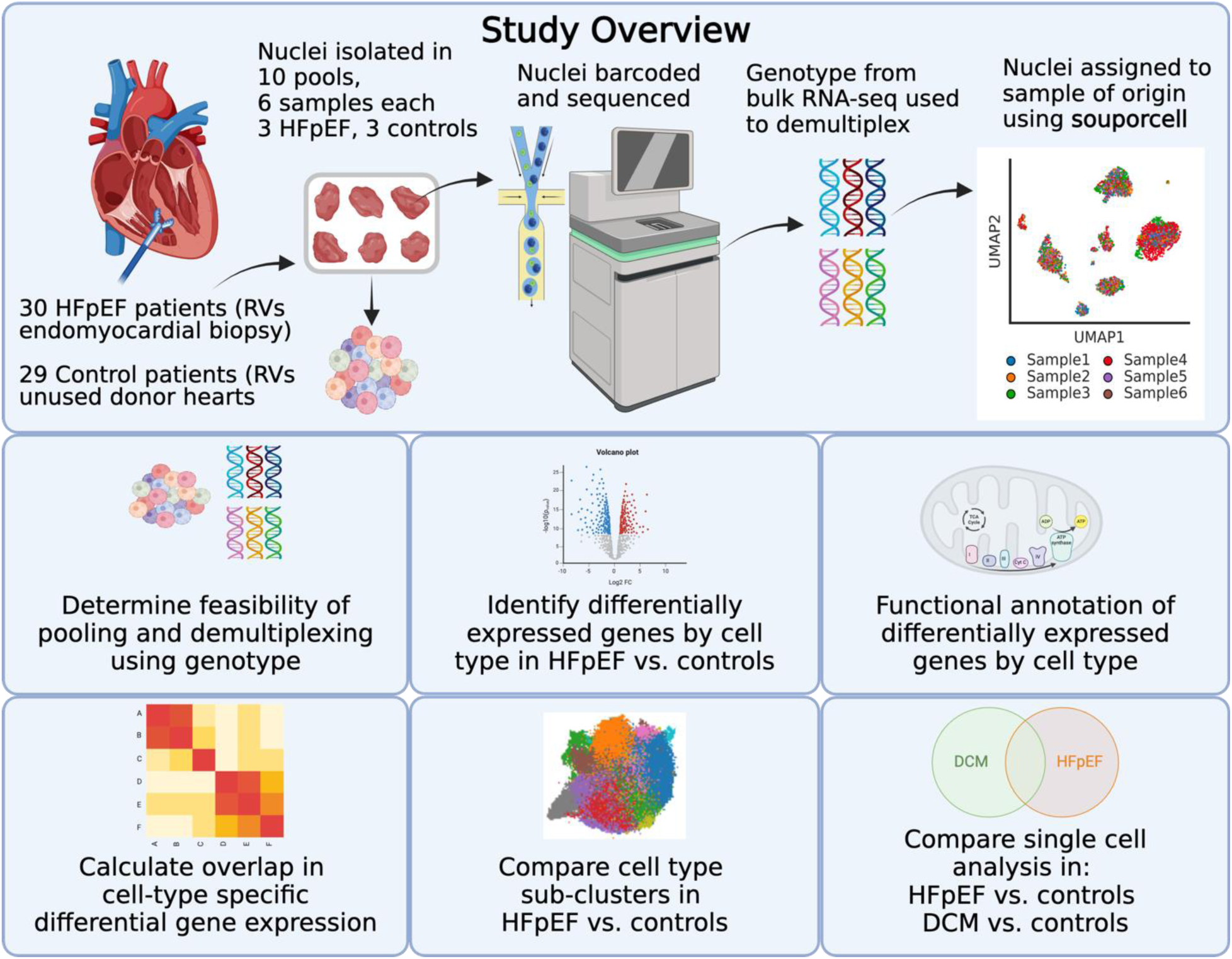
Study schematic and overview. Endomyocardial biopsies from patients with HFpEF were performed during clinically indicated right heart catheterization. Myocardial tissue samples were obtained from unused non-failing donor controls. Tissue samples were sorted into 10 pools with 3 HFpEF and 3 controls in each pool. Single nucleus RNA sequencing was performed. Nuclei were assigned to an individual sample based on genotype demultiplexing. Differential gene expression and functional annotation was performed in HFpEF vs controls. Overlap in differential gene expression was compared between cell types. Sub-clustering was performed in HFpEF vs controls. snRNA-seq in HFpEF vs controls was compared to DCM vs controls. RV, right ventricle; HFpEF, heart failure with reduced ejection fraction; DCM, dilated cardiomyopathy.

### Nuclei Isolation and Library preparation for single-nucleus RNA-sequencing

Nuclei were isolated using previously published protocols^14, 20^ with some modifications. Due to small biopsy size (2-3 mg), six were first pooled prior to nuclei isolation. Control samples were cut to match the size of the HFpEF biopsies so the proportion of control and HFpEF nuclei would be similar in each pool. The 6 samples (3 controls, 3 HFpEF) were embedded together in OCT and sectioned prior to nuclei isolation as described in the Supplement. Between 3,000 to 10,000 nuclei were loaded into a 10x Genomics Chip G and the 10x Genomics 3’ v 3.1 kit was carried out with slight modifications (Supplemental Methods). Library quality control (QC) and quantification were carried out using qPCR and Fragment analyzer, followed by sequencing on NovaSeq 6000, targeting ∼1 billion reads per library.

### Single-nucleus RNA-sequencing data processing, demultiplexing, and QC

snRNA-seq libraries were processed using CellRanger v4.0.0^21^. Specifically, BCL files were converted to FASTQs using the *mkfastq* pipeline. Prior to alignment, we trimmed homopolymer repeats and the template switch oligo sequence from R2 using cutadapt v1.18. ^22^ Resulting FASTQs were aligned and quantified to a pre-mRNA version of the GRCh38-2020-A transcriptome reference provided by 10X Genomics, using *cellranger count* with expected-cells 3000 to 10000. The resulting count matrix was processed with CellBender remove-background v0.2.0^23^ to identify non-empty droplets and remove ambient RNA and chimeric library fragments. A false positive rate (FPR) of 0.01 was used for all pools. We used scR-Invex (https://github.com/broadinstitute/scrinvex) to determine the proportion of reads that aligned exclusively to exons. To identify low quality libraries, we evaluated several CellRanger metrics (Supplemental Methods). Three libraries (Pools 1, 6, and 6b) were considered experimental failures and excluded from further analysis due to some combination of low valid barcodes, low rates of mapping confidently to the genome/transcriptome, low fraction of reads in CellRanger determined cells, and/or a low number of total genes detected (**OnlineExcelSupplement Table 1**, **Supplemental Figure 2**). The remaining 8 pools each contained 6 patients (3 HFpEF and 3 control). To assign nuclei to their patient of origin, we demultiplexed pooled libraries using *souporcell* v2.0^24^ (Supplemental Methods). Once nuclei were clustered based on genetic identity, we compared clusters to reference genotypes derived from bulk RNA-seq^9^ (Supplemental Methods). Five of the remaining 24 HFpEF samples were not detected in the genotype clusters used for demultiplexing, likely due to inadequate nuclei.

To control for inherent differences in QC metrics by cell type, we jointly aggregated 87,430 *souporcell* singlets to perform extensive nuclei QC as previously reported^14, 20^ and detailed in the Supplement. All single-cell analysis was performed using scanpy v1.9.1 unless otherwise stated. ^25^ We corrected for a batch effect for the patient of origin for each nucleus using Harmony v0.1.7 with default settings.^26^ To identify low quality nuclei, the following per-nucleus QC metrics were considered: 1) total number of unique molecular identifiers (UMIs), 2) total number of unique genes detected, 3) percent of UMI mapping to mitochondrial genes, 4) proportion of reads mapping exclusively to exons, 5) transcriptional entropy as estimated using the ndd library in Python (https://github.com/simomarsili/ndd/tree/master), and 6) doublet score as estimated in Scrublet v0.2.3^27^. After strict QC, 48,866 nuclei remained.

### Compositional testing and marker gene identification

We tested for differences in cellular composition between HFpEF and controls using *scCODA* v0.1.9 (Supplemental Methods). To identify marker genes for global cell types, we used a combination of the area under the receiver operating characteristic curve (AUC) calculated at the nucleus-level, and a formal differential expression test using a limma-voom^28, 29^ pipeline on pseudo-bulk aggregation (Supplemental Methods). Marker genes were selected as: 1) protein coding, 2) expressed in at least 25% of nuclei from the cell type of interest, 3) AUC > 0.7 in the nucleus-level test, and 4) logFC > 2 and adjusted p-value (Benjamini-Hochberg) < 0.01 based on the pseudo-bulk test.

### Patient-level principal component analysis (PCA)

PCA was performed using the *prcomp()* function in R 3.6.0. Gene counts were summed per patient across all cells or by cell type. After estimating dispersions with DESeq2, a variance stabilizing transformation was applied with the function *vst()*. The top principal components were then estimated based on the top 500 most highly variable genes. Details of this analysis are available in the Supplement.

### Differential expression between HFpEF and control

Differential gene expression between HFpEF and control samples was performed across all cell types (pseudo-bulk) and by cell type, similarly to Chaffin et al., 2022^14^ and Simonson et al., 2023^20^ and as detailed in the Supplement. Specifically, we generated pseudo-bulk expression matrices followed by differential expression testing using the limma-voom framework. Differential expression models were adjusted for sex and pool to control for batch effects. Multiple testing correction was performed within each cell type using Benjamini-Hochberg at false discovery rate (FDR)=0.05. Genes with a directionally consistent and significant (adjusted p-value < 0.05) association with both CellBender and CellRanger expression matrices, along with a background contamination heuristic ≤ 0.40, were considered significant. Estimates of logFC were compared to prior bulk RNA-seq in controls and HFpEF from the same biobanks, with overlapping samples in 19 HFpEF and 20 controls. ^9^

### Sub-clustering of major cell types

Sub-clustering of 6 major cell types (cardiomyocytes, fibroblasts, endothelial cells, mural cells, macrophages, and lymphocytes) was performed using single-cell Variational Inference (scVI) 0.17.3^30^ (Supplemental Methods). Marker genes for each sub-cluster were calculated as for the global map. Representative markers were selected as genes expressed in at least 15% of nuclei from the sub-cluster, with a logFC > 0.5 and FDR-adjusted p-value < 0.05 compared to all other sub-clusters of the cell type. To compare sub-clusters to previously reported populations, ^14, 15, 31–33^ we scored the transcriptional signatures from these previously published studies in our nuclei (Supplemental Methods).

### DCM and HFpEF combined analysis

snRNA-seq data from the 48,866 nuclei generated in this study were combined with 343,887 nuclei from 11 DCM and 15 controls previously reported, ^25^ mitigating batch effects between the studies (Supplemental Methods). Differential expression analysis by cell type was performed identifying genes in three categories: 1) differentially expressed between HFpEF (n_max_=19) and their respective controls (n_max_=24), 2) differentially expressed between DCM (n_max_=11) and their respective controls (n_max_=15), and 3) genes with a different effect in HFpEF versus control than DCM versus control. To achieve this, we used a similar pseudo-bulk testing approach as in the HFpEF versus control comparison where gene counts from all nuclei of a given cell type were summed within each patient. Using limma-voom, we fit a test for disease status adjusting for sex. We then extract three contrasts: HFpEF – Control_HFpEF_, 2) DCM-Control_DCM_, and 3) (HFpEF-Control_HFpEF_) – (DCM-Control_DCM_) to achieve the 3 comparisons of interest. We adjusted for multiple testing correction using the Benjamini-Hochberg correction at FDR=0.05. Differentially expressed genes unique to HFpEF were identified by being both different to controls and either unaltered or changed in the opposite direction in DCM. LogFC estimates from the combined DCM vs control (LV) and HFpEF vs control (RVS) snRNA-seq analysis were compared to the prior bulk RNA-seq data^9^ that had directly compared DCM vs HFpEF.

### Functional Annotation

Functional annotation was performed with Gene Set Enrichment Analysis of the Reactome database as described in the Supplemental Methods. A Jaccard similarity index was used to identify and remove redundant pathways as detailed in the Supplement. Genes associated with inherited cardiomyopathy or arrhythmias were identified based on their inclusion in the Invitae Arrhythmia and Cardiomyopathy Comprehensive panel (157 genes).

## RESULTS

### Study population

Of the 30 HFpEF patients and 29 controls initially included, 19 HFpEF patients and 24 controls remained for the final analysis. Eleven samples were excluded due to technical issues affecting two pooled samples (6 HFpEF, 5 controls) or unsuccessful genotype-based demultiplexing (5 HFpEF). Demographic and clinical characteristics of the final study population are presented in Table 1. Compared to controls, the HFpEF patients were older (median age 65 vs 58.5 years, p = 0.04), predominantly Black or African-American (84.2% vs 0%, p<0.0001, and had higher prevalence of hypertension (100% vs 58%, p=0.001) and diabetes (78.9% vs 12.5%, p < 0.0001). HFpEF subjects also had a higher body mass index (BMI, 43.2 vs 27.8kg/m^2^, p = 0.009) and lower estimated glomerular filtration rate (eGFR 35 vs 87mL/min/1.73 m^2^, p < 0.0001). The prevalence of coronary artery disease (5.3% vs 12.5%) and atrial fibrillation or flutter (15.8% vs 12.5%) was similar between the two groups. Resting invasive hemodynamic data for the HFpEF cohort (**Table 1**) was consistent with previously published data.^19^

**Table 1.** Clinical characteristics of study subjects.

| **Table 1. Clinical characteristics of study subjects.** |  |  |  |
| --- | --- | --- | --- |
|  | **HFpEF (N=19)** | **Normal (N=24)** | **p value** |
| **Subject Race/Ethnicity** |  |  | **< 1e-04** |
| Black or African-American, n (%) | 16 (84.2%) | 0 (0.0%) |  |
| White, non-Hispanic, n (%) | 3 (15.8) | 23 (95.8) |  |
| White, Hispanic, n (%) | 0 (0) | 1 (4.2) |  |
| **Age, years** | 65 (59.5, 70.0) | 58.5 (52.8, 63.2) | 0.036 |
| **Male Sex, n (%)** | 4 (21.1) | 12 (50.0) | 0.064 |
| **Medications** |  |  |  |
| ACEi/ARB, n (%) | 12 (63.2) | 7 (29.2) | 0.034 |
| Beta Blocker, n (%) | 12 (63.2) | 7 (29.2) | 0.034 |
| MRA, n (%) | 7 (36.8) | NA |  |
| Loop Diuretic, n (%) | 19 (100.0) | 0 (0.0) | < 1e-04 |
| **Past Medical History** |  |  |  |
| Hypertension, n (%) | 19 (100.0) | 14 (58.3) | 0.001 |
| Diabetes, n (%) | 15 (78.9) | 3 (12.5) | < 1e-04 |
| CAD, n (%) | 1 (5.3) | 3 (12.5) | 0.62 |
| AFib/Flutter, n (%) | 3 (15.8) | 3 (12.5) | 1 |
| **Clinical Characteristics** |  |  |  |
| BMI, kg/m<sup>2</sup> | 43.2 (37.1, 47.2) | 27.8 (23.5, 30.3) | < 1e-04 |
| SBP, mmHg | 141 (130.0, 155.0) | NA | NA |
| DBP, mmHg | 67 (64.0, 72.5) | NA | NA |
| **Echocardiography** |  |  |  |
| LVEF, % | 65 (55.0, 67.5) | 65 (60.0, 66.2) | 0.56 |
| LVEDD, cm | 4.2 (3.9, 4.8) | 4 (3.8, 4.5) | 0.21 |
| LVPW, cm | 1 (0.9, 1.3) | 0.9 (0.8, 1.1) | 0.17 |
| **Laboratory Data** |  |  |  |
| Creatinine, mg/dL | 1.6 (1.1, 1.9) | 0.9 (0.7, 1.3) | 0.024 |
| eGFR, mL/min/1.73m<sup>2</sup> | 35 (31.0, 57.0) | 87 (58.5, 103.5) | 0.009 |
| HbA1c, % | 6.6 (6.2, 9.1) | NA | NA |
| NTproBNP, pg/mL | 122 (67.5, 525.0) | NA | NA |
| **Invasive Hemodynamics** |  |  |  |
| RA, mmHg | 13 (9.5, 16.0) | NA | NA |
| PASP, mmHg | 45 (33.0, 56.0) | NA | NA |
| PADP, mmHg | 22 (19.0, 29.0) | NA | NA |
| PAm<sub>ean</sub>, mmHg | 31 (24.0, 35.5) | NA | NA |
| PAWP, mmHg | 20 (15.5, 25.5) | NA | NA |
| PA Saturation, % | 66.3 (63.7, 69.4) | NA | NA |
| CO, L/min | 5.9 (5.0, 6.4) | NA | NA |
| CI, L/min/m<sup>2</sup> | 2.5 (2.3, 2.9) | NA | NA |
| PVR, WU | 1.7 (1.2, 2.4) | NA | NA |
| Data presented as n (%) or median (25th-75th percentile). Fisher's exact test used for categorical variables. Mann-Whitney test used for continuous variables. CKD-EPI 2021 equation with creatinine used for eGFR calculations. HFpEF, heart failure with preserved ejection fraction; ACEi, angiotensin converting enzyme inhibitor; ARB, angiotensin II receptor blocker; MRA, mineralocorticoid receptor antagonist; CAD, coronary artery disease; AFib, atrial fibrillation; BMI, body mass index; SBP, systolic blood pressure; DBP, diastolic blood pressure; LVEF, left ventricular ejection fraction; LVEDD, left ventricular end diastolic diameter; LVPW, left ventricular posterior wall; eGFR, estimated glomerular filtration rate; HbA1c, hemoglobin A1C; NTproBNP, N-terminal pro-B type natriuretic peptide; RA, right atrial pressure; PASP, pulmonary artery systolic pressure; PADP, pulmonary artery diastolic pressure; PAm<sub>ean</sub>, mean pulmonary artery pressure; PAWP, pulmonary artery wedge pressure; PA, pulmonary artery; CO, cardiac output; CI, cardiac index; PVR, pulmonary vascular resistance; wu, Wood units; N/A, data not available for controls. |  |  |  |

### Single nucleus UMAP generation demonstrates feasibility of genotype-based demultiplexing

The feasibility of pooling samples and demultiplexing using genotype information from bulk RNA-seq is shown in **Supplemental Figure 1**. Using samples from the same 6 individuals, we identified the same number and type of cells in both the pooled and individually run samples. Cell composition was similar and logFC estimates had strong correlation between the two techniques. Based on this, we proceeded with the pooling strategy. After exclusion of 2 pools based on QC (**Supplemental Table 1, Supplemental Figure 2**), 119,226 CellBender-determined non-empty droplets were demultiplexed by *souporcell* using genotype information from bulk RNA-seq. Seventeen nuclei clusters were observed including 5 clusters of doublet/unassigned nuclei (**Figure 2a**), with overlap in the pools represented in each cluster (**Figure 2b**). The *souporcell*-derived assignment of genotype clusters to an individual patient ID matched only the list of patient IDs that were included in each pool (**Figure 2c**), and demultiplexing had a high concordance between the known sex and inferred sex based on gene expression (**Supplemental Figure 3**). There were 4 pools with a genotype cluster that did not map to any individual (unassigned), potentially due to small biopsy size. Doublet nuclei had 2 genotypes identified within a droplet. Overall, 76.7% of nuclei were assigned to an individual and this varied slightly by cell type and pool (**Figure 2d, 2e, Supplemental Table 2**). A total of 87,430 nuclei were successfully demultiplexed and assigned to an individual sample.

**Figure 2.**
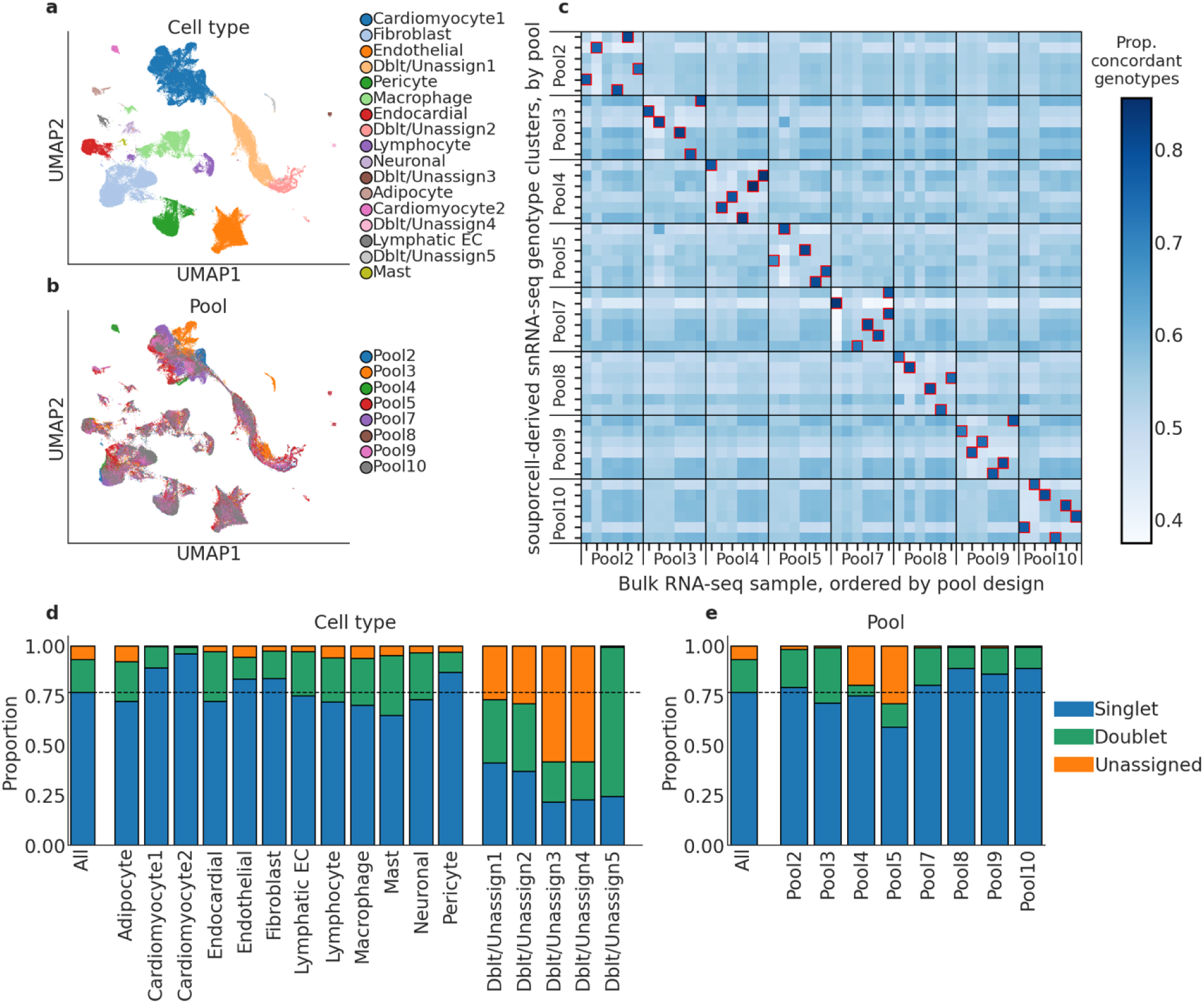
**Demultiplexing biopsy nuclei from snRNA-seq pools using genetics. a-b**, UMAP embedding of all CellBender-determined non-empty droplets (n=119,226) colored by cell type as identified with Leiden clustering (**a**) and pool of origin (**b**). c, Heatmap displaying the proportion of concordant genotypes for each *souporcell* derived genotype cluster in each pool (k=6, y-axis) and each bulk RNA-seq sample (x-axis). Bulk RNA-seq samples are ordered by the pool in which they were included for snRNA-seq data generation. *souporcell* genotype clusters assigned to an RNA-seq sample based on a high concordance between *souporcell* genotypes and bulk RNA-seq genotypes are outlined in red. **d, e,** Proportion of nuclei assigned as singlets, doublets, and unassigned by cell type as identified with Leiden clustering (**d**) and pool (**e**). Dblt, Doublet; Unassign, Unassigned.

Next, extensive QC was performed to remove doublets, cytoplasmic contamination, and technical failures. Pre-QC uniform manifold approximation and projection (UMAP) representations are shown in **Supplemental Figure 4**, with subsequent UMAP representation highlighting the nuclei removed from the initial QC (37,165) and the subsequent subcluster cleaning (1399). After QC, 48,866 nuclei remained from 19 HFpEF and 24 controls. Identified marker genes for each nuclei cluster (**Supplemental Table 3, Supplemental Figure 5**) were used to annotate cell-types in the final UMAP (**Figure 3a**). Two distinct endothelial cell clusters were identified, subsequently labeled endothelial 1 (ET1, the more abundant cluster) and endothelial 2 (ET2). The most abundant cell types were cardiomyocytes, representing on average 26.6% of nuclei in each patient, followed by fibroblasts, ET1, pericytes, and macrophages representing on average 24.1%, 16.7%, 10.4%, and 5.5% of nuclei (**Figure 3b**). While proportions of cell types varied among individuals (**Supplemental Figure 6**), similar cellular composition was found in HFpEF and controls for all types but endocardial cells (**Supplemental Table 4, Figure 3b**).

**Figure 3.**
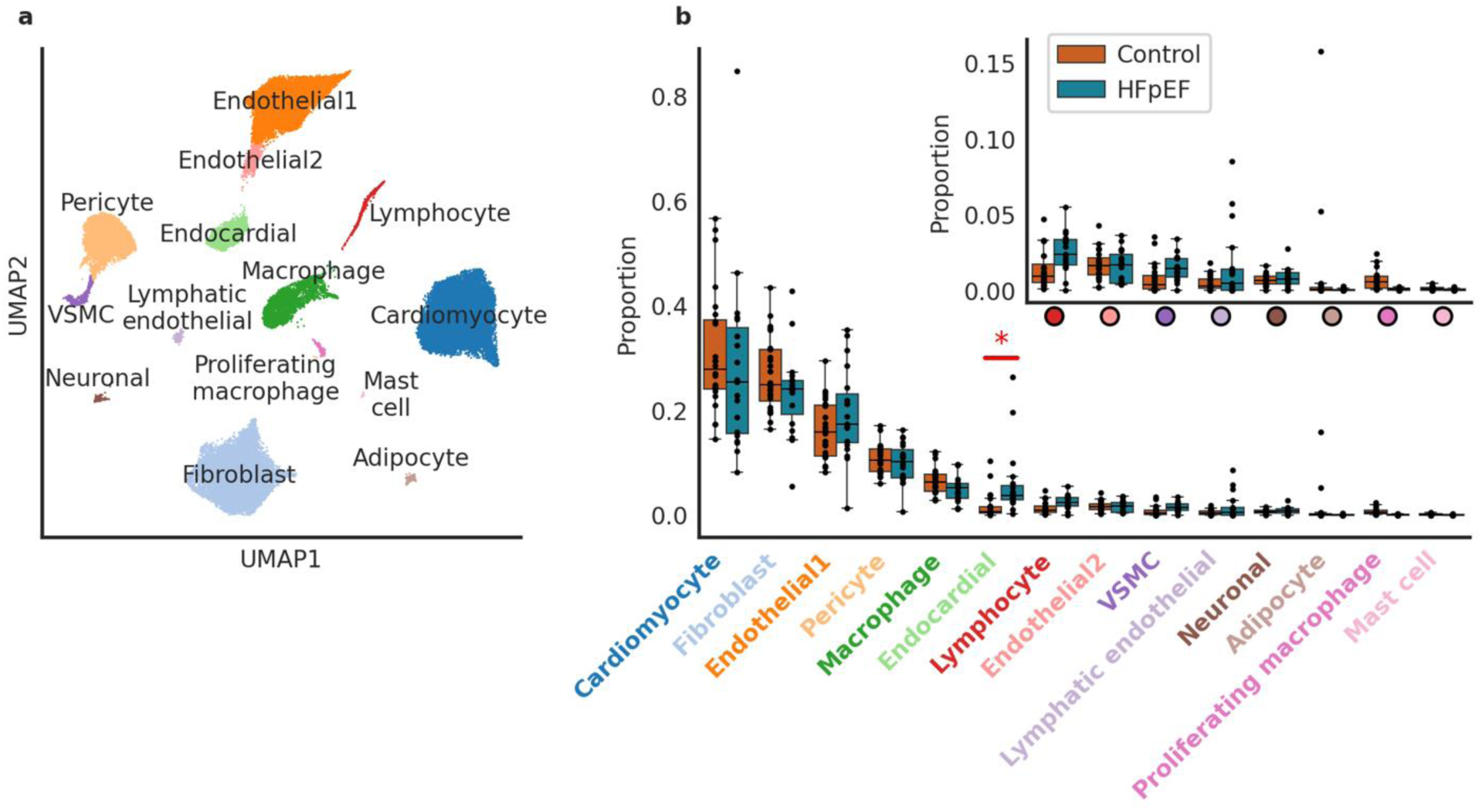
Aggregation of 48,866 nuclei in a comprehensive single-nucleus map. **a)** UMAP embedding of 48,866 nuclei derived from 8 pools across 43 patients. Nuclei are colored by cell type. **b)** Box plots demonstrating the compositional differences between HFpEF cases and controls. Cell types are colored as in panel **a** and inset depicts the composition of rarer cell types for improved viewability. Significant compositional shifts detected with *scCODA* are denoted with an *. Center line, median; Box limits, upper and lower quartiles; Whiskers, 1.5x interquartile range.

### Differential gene expression between HFpEF and controls reveals substantial transcriptional dysregulation across cell types

Nine of the 14 identified cell-types had sufficient nuclei in both HFpEF and controls to perform differential expression analysis. Patient level principal component analysis (PCA) demonstrated separation of HFpEF vs controls for each cell type (**Supplemental Figure 7**). Of the 14,368 genes identified in cardiomyocytes in both patient groups, 36% (5159) were differentially expressed; nearly half (49%) of 12,166 genes were differentially expressed in fibroblasts (**Figure 4a, Supplemental Table 5**). Endothelial cells (ET1 cluster, 2143 of 6271 [25%]), pericytes (1812 of 5571 [25%]), and macrophages (1405 of 4756 [23%]) also displayed substantial differential gene expression, whereas endocardial cells, lymphocytes, VSMCs and a second endothelial cell cluster (ET2) had <10% detected genes differentially expressed (**Figure 4a, Supplemental Table 5**). Volcano plots with annotation of the top 8 up- and down-regulated genes are displayed in **Figure 4b**. Estimated logFC between HFpEF and control based on the pseudo-bulk data for all cell types significantly correlated with the prior bulk RNA-seq data ^9^ (r = 0.68, p< 2.2e-16, **Supplemental Figure 8**). We further examined for overlap between genes differentially expressed for each cell type and those identified in pseudo-bulk analysis to determine how well each cell type was represented by bulk RNA-seq approaches (**Supplemental Figure 9**). Most DEG in cardiomyocytes (72.0%) overlapped with pseudo-bulk, whereas this was less for lower abundant cell types (macrophage, 52.2%; VSMC, 50.0%; lymphocyte, 41.7%). This highlights the ability of snRNA-seq to identify differentially expressed genes in lower abundant cell types that could be missed by bulk RNA-seq.

**Figure 4.**
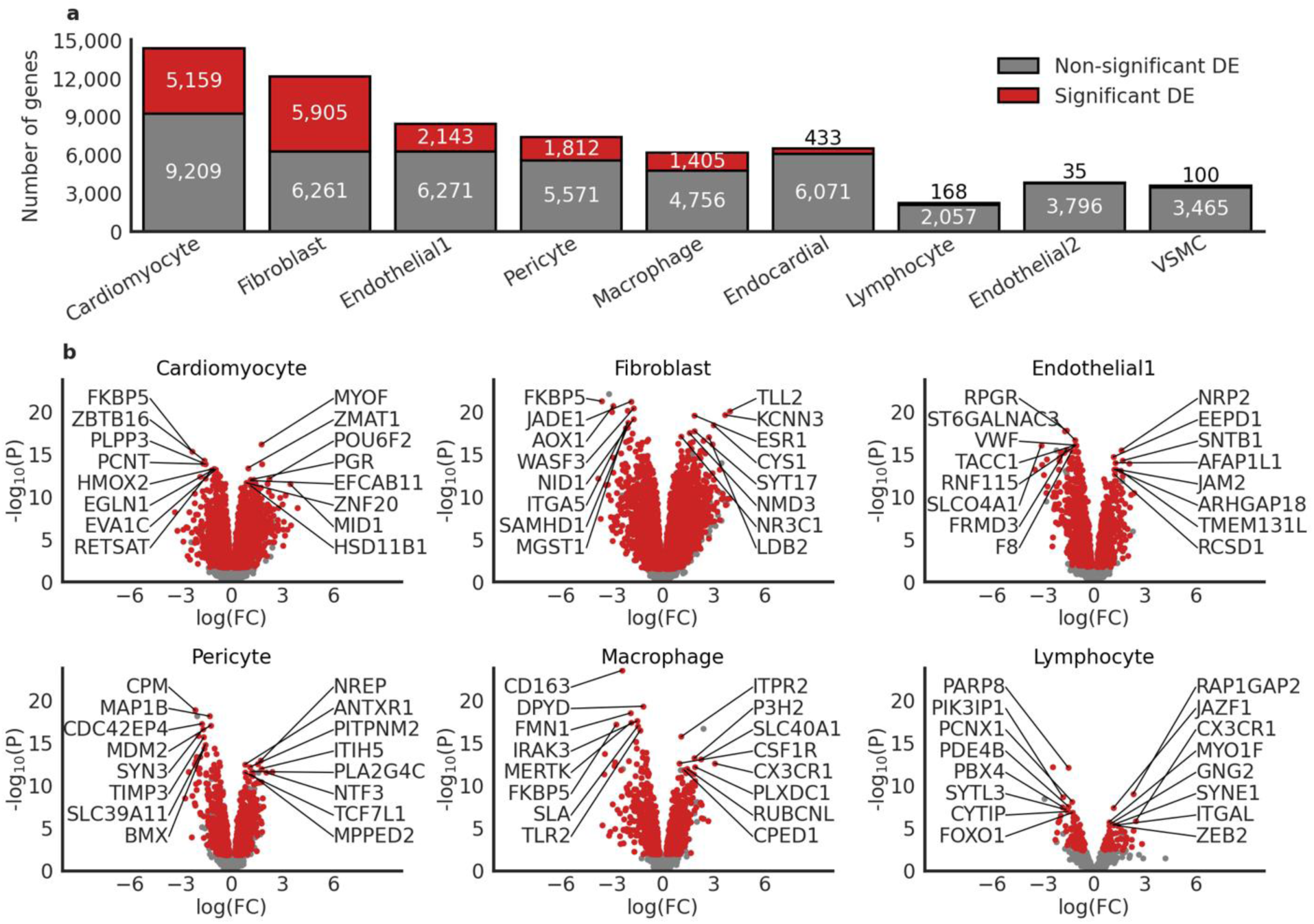
Differential expression between HFpEF and control at single-cell resolution. **a)** Bar plots showing the number of significant (red) and non-significant (gray) genes identified in the differential expression (DE) testing by cell type. **b)** Volcano plots for differential expression between HFpEF and control in six cell types. The x-axis represents log2 fold-change (logFC) estimates between HFpEF compared to control and the y-axis is the -log10(P-value), based on the CellBender primary analysis. Dots colored red represent significantly differentially expressed genes, defined as having a low background probability heuristic (<0.40), significant differential expression with both CellBender and CellRanger counts (FDR < 0.05), and a concordant effect estimate with both CellBender and CellRanger. The top 8 up- and down-regulated protein coding genes are labeled.

### Functional annotation in HFpEF vs controls identifies shared and unique pathways among the cell types

Functional annotation was performed by gene set enrichment analysis (GSEA) to identify enriched Reactome pathways in cardiomyocytes, endocardial cells, endothelial1, fibroblasts, macrophages, pericytes and vascular smooth muscle cells (VSMCs) (**Figure 5, Supplemental Table 6**). After curation to avoid redundancy, we identified several pathways shared by more than one cell type. Most had a negative normalized enrichment score, signifying enrichment of downregulated genes. Cardiomyocytes and fibroblasts had the highest number of overlapping pathways (**Figure 5a**). Highlights of the overlapping pathways included broad categories related to translation, immune system activation, metabolism, protein quality control, and metabolism of RNA and transcription. Reactome pathways unique to each cell type are shown in **Figure 5b**. Highlights of enriched pathways in cardiomyocytes include metabolism (lipoprotein metabolism, PP2A dephosphorylation of metabolic factors, beta-oxidation, respiratory electron transport, pyruvate metabolism and TCA cycle), extracellular matrix organization, calcium handling, and growth factor signaling. Highlights in fibroblasts include regulation of gene expression, mTORC and nitric oxide signaling, and metabolism of amino acids and glucose. To gain further insight into cell-specific differences in pathways, we selected the pathway with the lowest percentage of genes shared across core enrichments for visualization as a heatmap. The adaptive immune system pathway was enriched in cardiomyocytes and macrophages, yet most core enrichment genes in cardiomyocytes were down-regulated and distinct from the core enrichment genes in macrophages that were up-regulated (**Supplemental Figure 10).** For further functional annotation specific to genes related to inherited cardiomyopathies and arrhythmias, we examined the DEG that overlapped with 157 genes in the Invitae genetic testing panels (**Supplemental Figure 11**). A substantial number of genes were significantly downregulated in cardiomyocytes (38/157 genes) and other cell types.

**Figure 5.**
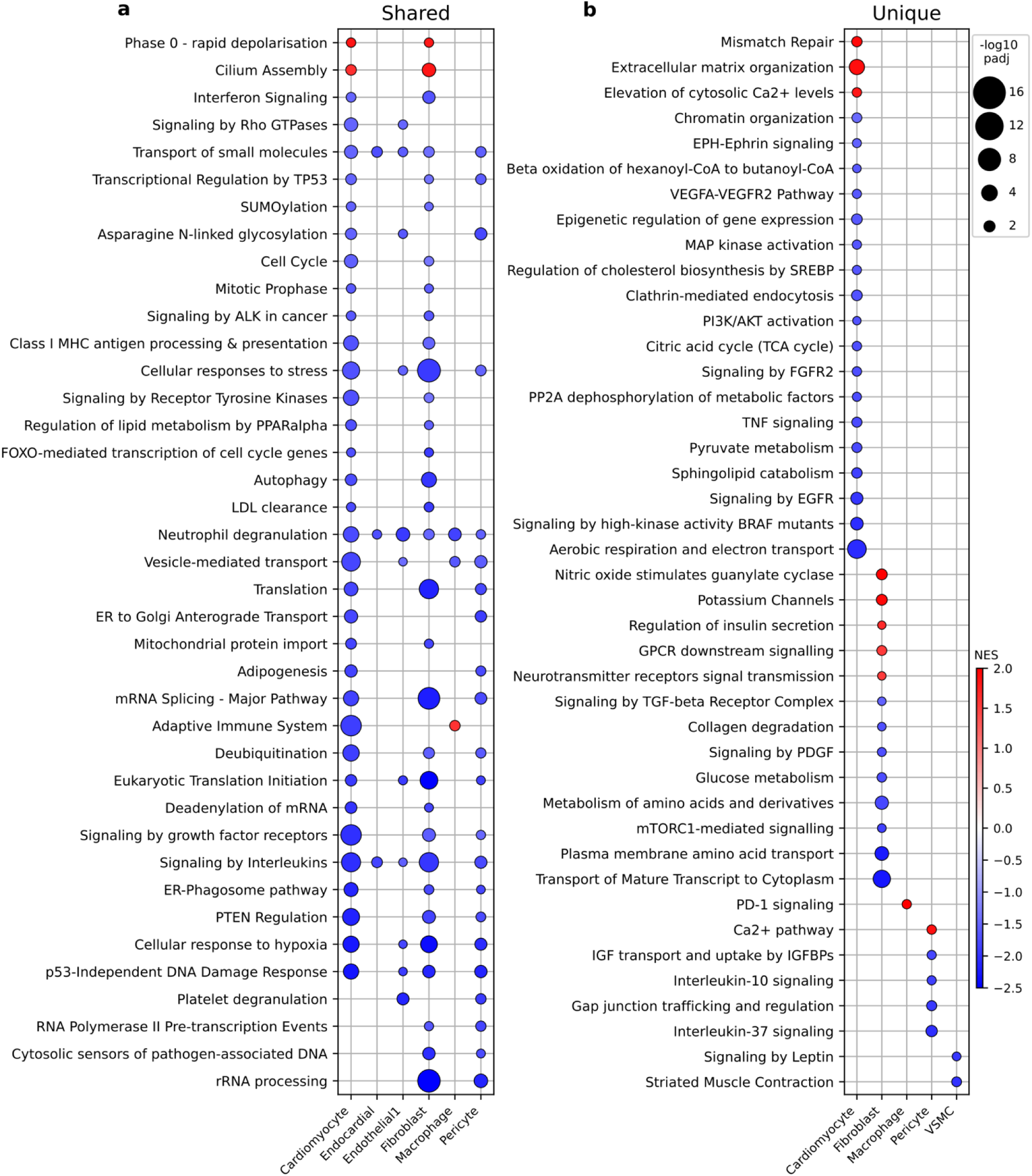
Gene Set Enrichment Analysis of differentially expressed genes in HFpEF vs controls. Dot plots for Reactome pathways significantly enriched by differentially expressed genes in HFpEF vs controls in **(a)** multiple cell types and **(b)**, one cell type. Pathways were curated using a Jaccard similarity index to identify redundant pathways. Pathway names were shortened for visibility. Cell types are displayed on the x-axis (CM, cardiomyocyte; EC, endocardial; ET1, endothelial 1; Fibro, fibroblast; Macro, macrophage; Peri, pericyte; VSMC, vascular smooth muscle cell). Dot size corresponds to the -log10 of the adjust p-value for each GSEA pathway, with larger dots indicating a smaller p-value. P values were adjusted for multiple comparisons using the Benjamini-Hochberg method. Dot color corresponds to normalized enrichment score (NES) for each pathway.

### Sub-clustering reveals the absence of activated fibroblasts in HFpEF

Sub-clustering within the major cell types identified sub-types for cardiomyocytes (n=7), fibroblasts (n=5), endothelial cells (n=10), mural cells (pericytes and VSMC, n=6), macrophages (n=6), and lymphocytes (n=3) (**Figure 6, Supplemental Figures 12-16**). Genes specific to each sub-cluster are provided in **Supplemental Table 7**. The transcriptional signatures of these sub-populations were further compared to cell states signatures reported in prior single cell RNA-seq studies (**Supplemental Figures 17-22**). Notably, among the 5 fibroblast sub-clusters, FB-0 was more prevalent in controls, while FB-1 was more prevalent in HFpEF, likely representing global changes in gene expression (**Figure 5c**). Marker genes for FB-4 were consistent with those of activated fibroblasts (**Figure 5d, Supplemental Figure 17**), yet were not identified in HFpEF samples and derived from a few control samples (**Figure 5c**).

**Figure 6.**
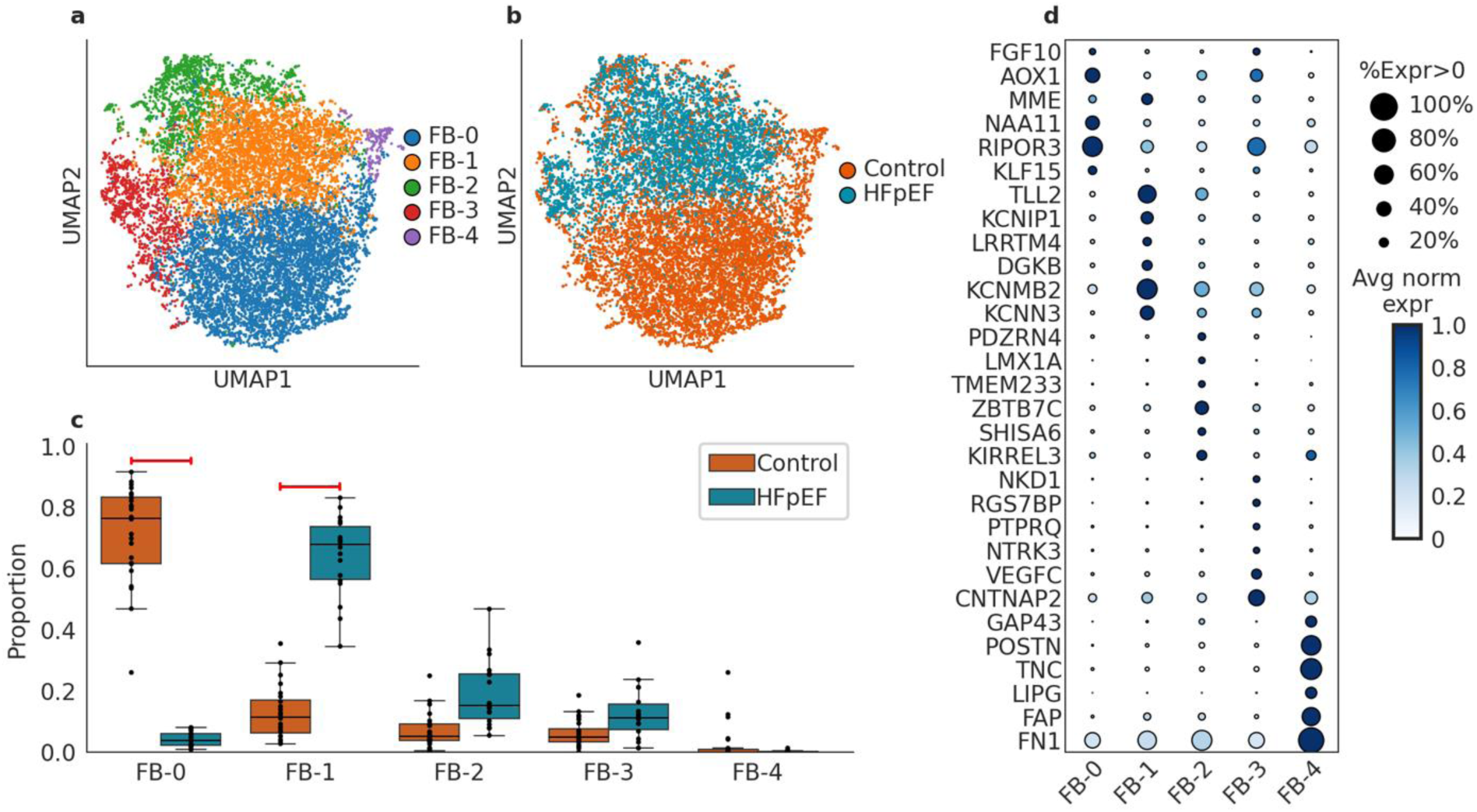
Sub-populations of fibroblasts. **a)** UMAP embedding of 12,562 fibroblasts based on scVI batch-corrected latent space colored by Leiden sub-clusters at resolution 0.3. **b)** UMAP embedding of fibroblasts colored by disease status. **c)** Box plots showing the proportion of each fibroblast sub-cluster among all fibroblasts in patients, separated by HFpEF and control. Center line, median; Box limits, upper and lower quartiles; Whiskers, 1.5x interquartile range. **d)** Dot plot showing the expression of the top marker genes for each fibroblast sub-cluster. The size of each dot represents the percent of nuclei that express the gene at non-zero levels (%Expr>0) and the shade represents the average log-normalized expression, scaled to the maximum value across sub-clusters for each gene (Avg norm expr).

Within other cell types, credible shifts in composition in HFpEF often seemed to reflect overall changes in transcription rather than distinct cell states (e.g., CM-0, ET-0, ET-2, ET-4, ET-6, Peri-0, Peri-1, Peri-2, VSMC-1, MP-0, and MP-2). However, some distinct cell states displayed shifts in HFpEF, including an increase in endocardial cells (EC-1, consistent with the global analysis), an increase in monocytes (MP-4), and a decrease in one population of T-cells (LC-0). Additionally, we were able to identify populations of arterial endothelial cells (ET-1), venous endothelial cells (ET-5), lymphatic endothelial cells (LEC), cycling or proliferating macrophages (MP-3), B-cells (MP-5), and NK cells (LC-1), but none of these sub-clusters showed a significant shift in composition in HFpEF.

### Comparison of HFpEF to DCM reveals most divergent transcriptional changes occur in cardiomyocytes

To understand the transcriptional differences between HFpEF and DCM, we next combined the HFpEF and NF control snRNA-seq with prior snRNA-seq in DCM and NF controls generated from LV tissue. Though the two experiments studied different tissue sources (RVS or LVS), our prior study showed gene expression in RVS vs LVS were highly correlated in both DCM and controls.^9^ Therefore we proceeded with integrating the two experiments to compare DCM and HFpEF. The combined UMAP identified typical cell types in myocardial biopsy tissue with mixing of the disease groups within each cell type (**Supplemental Figure 23**). Patient level PCA demonstrated significant overlap between the controls from both experiments suggesting comparability between datasets. Deviations between HFpEF and DCM were observed in specific cell types, particularly cardiomyocytes **(Figure 7a)**. Cardiomyocytes had the highest number of genes (1482 genes) with significantly different logFC comparing HFpEF vs control to DCM vs control (e.g. different between HFpEF vs DCM), while other cell types had few genes significantly different between HFpEF and DCM (**Figure 7b, Supplemental Table 8)**. A subset of genes that were only differentially expressed in HFpEF or were differentially expressed but in opposite directions between HFpEF and DCM (vs controls) are highlighted in **Figure 7c** (cardiomyocytes) and **Supplemental Figure 24** (non-cardiomyocytes). These genes had an absolute logFC of at least 1 and a minimal threshold of expression for the reference group. Of the DE genes between HFpEF and DCM in the prior bulk RNA-seq, the snRNA-seq logFC estimates in pseudo-bulk and cardiomyocytes moderately correlated with the prior bulk RNA-seq, while other cell types had only a weak correlation (**Supplemental Table 9**). These data suggest the differences between HFpEF and DCM in the bulk RNA-seq were most representative of DE in cardiomyocytes.

**Figure 7.**
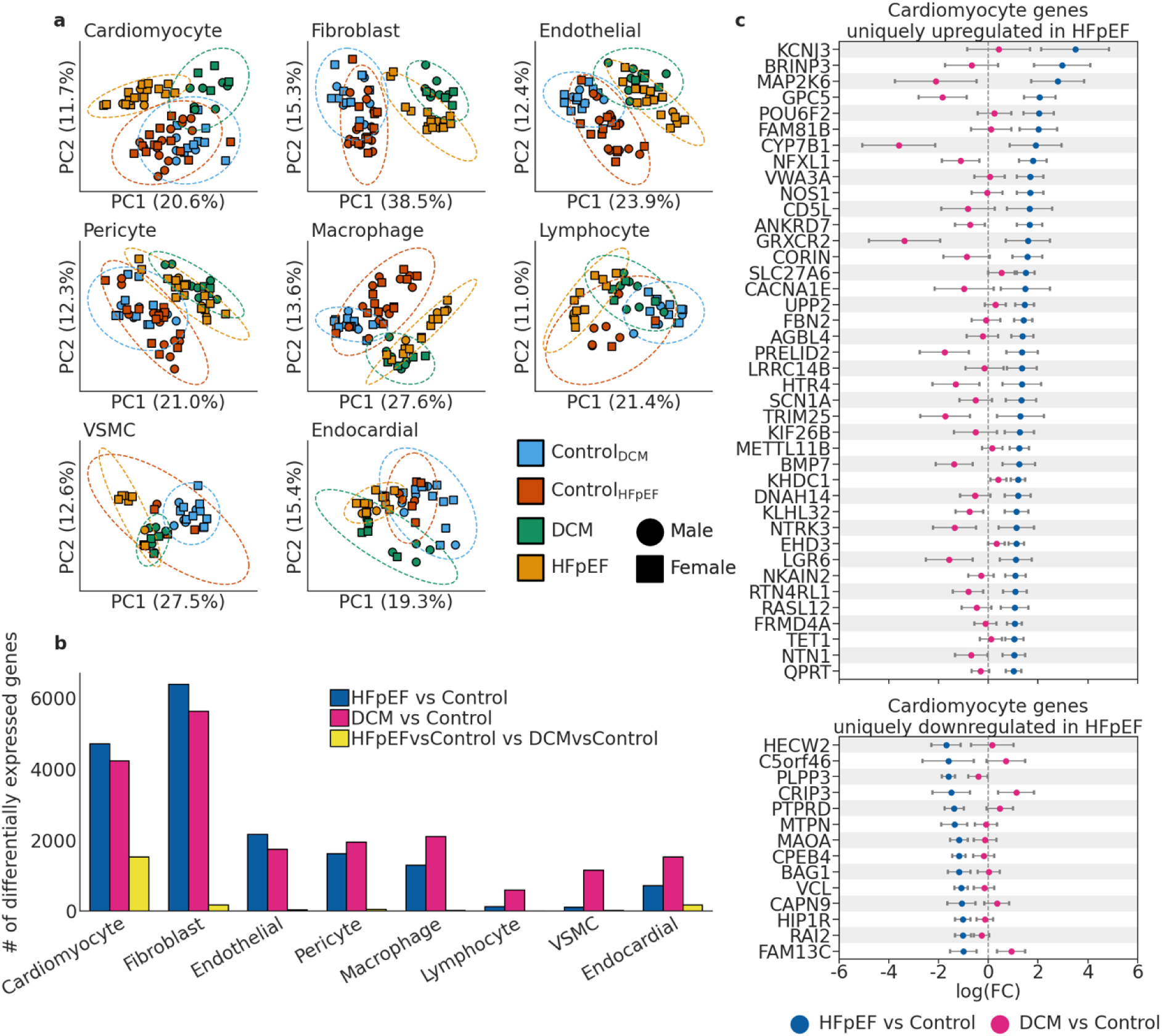
Transcriptional differences between HFpEF and DCM. **a)** Principal component (PC) analysis by cell type comparing broad transcriptional differences between controls, heart failure with preserved ejection fraction (HFpEF), and dilated cardiomyopathy (DCM) patients. The percent of variation explained by each PC is shown on the respective axis. Normal-probability contours based on a 95% data ellipse are shown for each group. **b)** Bar plots showing the number of genes significantly differentially expressed by cell type in the comparison of HFpEF to control, DCM to control, and showing a significantly different effect in HFpEF versus control than DCM versus control (HFpEFvsControl vs DCMvsControl). **c)** Forest plots display logFC in cardiomyocytes for selected genes uniquely down- or up-regulated in HFpEFvsControl, and either unchanged or differentially expressed in the opposite direction in DCMvsControl. LogFC is from the HFpEFvsControl comparison for HFpEF, and the DCMvsControl comparison for DCM. Error bars represent 95% confidence intervals. Genes selected for visualization had 1) absolute log_2_ fold change (logFC) ≥ 1, 2) Benjamini-Hochberg corrected p-value < 0.05, 3) proportion of nuclei with detectable expression of that gene at least 0.1 in HFpEF for upregulated genes, or at least 0.1 in controls for downregulated genes, and 4) were protein coding. LogFC was calculated using CellBender counts. VSMC, vascular smooth muscle cell.

## DISCUSSION

To our knowledge, the present study provides the first snRNA-seq analysis of myocardium comparing HFpEF to controls and further integrates the snRNA-seq data in DCM vs controls. Our study yields several key findings. First, we demonstrated the feasibility of snRNA-seq on small human heart biopsies by pooling samples to increase nuclei yield and subsequently using genotype-based demultiplexing. Second, we identified numerous differentially expressed genes in HFpEF compared to controls across cardiomyocytes, fibroblasts, endothelial cells, pericytes, and macrophages. Third, functional analysis revealed shared enrichment of pathways related to transcription/translation, immune activation, metabolism, and protein quality control across multiple cell types, consistent with previous tissue-based omics studies in HFpEF. Fourth, unlike DCM, sub-clustering did not reveal an activated fibroblast population in HFpEF. Finally, we show that cardiomyocytes exhibit the greatest transcriptional divergence between HFpEF and DCM, while other cell types show fewer differences. These results illuminate potential shared and cell-type-specific pathogenic mechanisms in HFpEF.

The strategy to pool tissues and use genotype-based demultiplexing was used since individually the biopsies did not yield sufficient nuclei for the 10X platform to work effectively due to their small size. That the pseudo-bulk data derived from snRNA-seq analysis strongly correlated with bulk RNA-seq data from the identical patients further supports this approach. By pooling groups of controls and HFpEF samples to the same shared background ambient RNA, this allowed for higher throughput at lower cost. While technical effects may vary between libraries, we could control for these differences with a fixed effect for pool in the differential expression model as both HFpEF and control samples were represented in each pool.

The snRNA-seq analysis identified 14 clusters with similar cell type identifications and proportions in other snRNA-seq analyses in human myocardium. ^14, 16–18, 33^ Several of these prior studies found a lower proportion of cardiomyocytes in DCM and HCM compared to controls, ^14,15, 33^ despite there being greater LV mass in both disease groups. This could reflect myocyte hypertrophy while other cell types proliferated, lowering the proportion of myocytes, or could be due to cardiomyocyte loss. However, we did not find a similar overall change in HFpEF, and this could reflect less hypertrophic remodeling, less transcriptional complexity in non-myocytes making it more difficult to demultiplex pooled data, or differences in tissue sampling between control and HFpEF hearts. The HFpEF samples had a higher proportion of endocardial cells which might reflect use of endomyocardial biopsies. While control tissue came from the RVS, it was aliquoted from larger pieces making it hard to precisely replicate the composition.

Similar to data from DCM and HCM, we found the largest number of differentially expressed genes in cardiomyocytes and fibroblasts. ^14^ Consistent with the myocardial proteome in HFpEF,^11^ downregulated genes were enriched in pathways related to transcription and translation, protein quality control, and metabolism. Interestingly, metabolic pathways were broadly downregulated in multiple cell types, suggesting general factors such as obesity, diabetes, hemodynamic and neurohormonal stressors are drivers for such changes. Prior HFpEF studies found evidence of impaired fatty acid and glucose utilization by the myocardium ^9–11, 34^, including a study assessing arterial versus coronary sinus metabolites to determine net cardiac uptake. ^35^ Similar patterns have been reported in DCM, which like HFpEF, has depressed fatty acid metabolism genes in multiple cell types ^14, 15^. Interestingly, this is found in DCM where obesity/diabetes is less marked than in HFpEF, and where it improves with mechanical unloading. ^18^ While genes related to oxidative phosphorylation were previously found upregulated in HFpEF myocardium, ^9^ in the current analysis they were downregulated in cardiomyocytes specifically. This could relate to differences inherent to nuclear vs cytosolic transcriptomics which have previously shown to bear discrepancies in oxidative phosphorylation genes. ^36^

Proteomic analyses of peripheral blood in HFpEF have identified higher levels of inflammatory proteins,^20–22^ and histologic studies suggested an increase in macrophage infiltration in the heart in HFpEF.^19, 37^ Although macrophages in general did not differ in abundance between HFpEF and controls in our study, one sub-cluster identified as monocytes were more abundant in HFpEF. Prior bulk RNA-seq analysis of HFpEF vs controls ^9^ did not identify inflammation or immune activation as an enriched pathway. Here, several pathways related to inflammation and immune activation were enriched in multiple cell types, yet they were generally down-regulated in non-immune cells, as has been reported in DCM. ^14^ Several of the pathways had many genes ubiquitously expressed in other cell types, and these could serve non-immune/inflammatory functions in these cells. This was most pronounced in the adaptive immune system pathway, which was enriched in cardiomyocytes and macrophages, but in opposite directions.

Of the genes downregulated in cardiomyocytes and unchanged in macrophages in the adaptive immune system pathway, several related to protein quality control (*KLHL3, PSMB7, PSMC4, PSMA7, TRIM63, SEC23A, KBTBD8, PPIA*), cellular signaling (*RLIM, PIK3R1, TRAF7, RAF1, PRKG1, PPP2CA, IKBKG*), and cytoskeletal components (*VASP, TUBB1, TUBA3D*). Genes downregulated in cardiomyocytes and upregulated in macrophages also included some related to protein quality control (*RNF220, CBLB, UBA3, SH3RF1, RNF19A, RNF14, UBE2W, WWP1*), cellular adhesion (*ITGAV*), fatty acid metabolism (*CD36*), and cellular signaling (*PTPRJ, MAPKAP1, SOS1, PPP2R5A*). Genes upregulated in only macrophages were specific to immune cells, including major histocompatibility proteins and their receptors (*CD74*), and others related to immune cell signaling (*ITPR2, BLNK, ZNRF2LYN, LILRB1, PRKCB, TAB2, VAV1, PTPRC, RAP1GAP2, PLCG2, PAG1, FYB1*) and cellular adhesion/migration (*ITGA4, EVL*). These findings highlight the value of single cell analyses to uncover discordant transcriptomic profiles, particularly in lower abundant cell types. The macrophage gene signature may have been diluted in the prior bulk RNA-seq experiment, limiting our ability to identify immune activation as an enriched pathway.

Contrary to prior studies in DCM, ^14^ we did not find a higher proportion of activated fibroblasts in HFpEF vs controls. However, there were many differences in cardiomyocytes from HFpEF vs DCM. Among cardiomyocyte DEGs upregulated in HFpEF and reduced in DCM is *MAP2K6* (or *MAPKK6*), which is activated mitogen activated kinase p38. Intriguingly mice with *MAPKK6* cardiac overexpression develop marked diastolic dysfunction without chamber dilation^38^. Another is tripartite motif containing 25 (*TRIM25*), a ubiquitin E3 ligase that regulates gene expression by binding to a distinct set of coding and non-coding RNAs^39,40^ and microRNAs^41^. Overexpression of *TRIM25* increases macrophage inflammatory polarization, while knockdown reduces inflammation, and collagen deposition and fibrosis^42^. We also found *NOS1* uniquely upregulated in HFpEF. *NOS1* is a key component of a now common HFpEF mouse model combining high fat diet with inhibition of nitric oxide synthase.^43^ Among genes specifically downregulated in HFpEF were the phospholipid phosphatase *PLPP3* which has pleotropic effects on lipid signaling and plays a role in cardiac hypertrophy and decreased mitochondrial bioenergetics^44^. Others include protein tyrosine phosphatase receptor type D (*PTPRD*), which activates *PPAR* gamma 2 and regulates insulin signaling^45^. These and others listed in Figure 7c may identify novel mechanistic and therapeutic targets.

This study has several limitations. The HFpEF and control groups differed in demographics, most notably in self-identified race. The HFpEF group was predominantly Black/African-American, reflecting the patient population served by Johns Hopkins Hospital, a diverse, urban community. In contrast, the donor heart population, from which the controls were derived, has a much lower percentage of Black/African-American individuals. Additionally, controls were younger and less obese. Tissue harvesting methods also differed. While control tissue was derived from the same endocardial region of the right ventricular septum as the HFpEF samples, the control and DCM samples used in prior studies were obtained from whole excised hearts after anterograde perfusion with cold cardioplegia. This procedure was not feasible for the HFpEF subjects. Pooling samples may have reduced the recovery of less transcriptionally complex cell types, as genetic demultiplexing relies on reads from the snRNA-seq itself. In rare instances, discrepancies arose between our snRNA-seq findings and bulk RNA-seq data, potentially due to differences between nuclear and cytoplasmic RNA isolation. Finally, the prior DCM vs. control experiment used in our combined analysis yielded a much higher number of nuclei, especially for less abundant cell types such as lymphocytes, which may have limited our power to detect differences between DCM and HFpEF.

In summary, our study demonstrates the feasibility of snRNA-seq on small myocardial biopsies using genotype-based demultiplexing, yielding several significant insights. We observed broad downregulation of metabolism, translation, and protein quality control pathways across multiple cell types in HFpEF myocardium. While non-cardiomyocyte cell populations in HFpEF exhibit transcriptomic profiles similar to those in DCM, cardiomyocytes show distinct gene expression patterns in HFpEF compared to DCM. These HFpEF-specific cardiomyocyte genes warrant further investigation.

## ACKNOWLEDGEMENTS

We thank the patients, donors, and donor families who agreed to participate in the study. We thank the Gift-of-Life Donor Program, Philadelphia, PA, who helped provide non-failing heart tissue from unused donor hearts for this study.

## NON-STANDARD ABBREVIATIONS AND ACRONYMS

CM: cardiomyocytes
DCM: dilated cardiomyopathy
GSEA: gene set enrichment analysis
HFpEF: heart failure with preserved ejection fraction
logFC: log-2 fold change
LVS: left ventricular septum
OCT: optimal cutting temperature
PCA: principal component analysis
qPCR: quantitative polymerase chain reaction
RVS: right ventricular septum
snRNA-seq: single nucleus RNA sequencing
TCA: tricarboxylic acid
UMI: unique molecular identifier
VSMC: vascular smooth muscle cells

